# Lipid Changes due to Low O-GlcNAc and the Downstream Consequences

**DOI:** 10.64898/2026.10.13.758586

**Authors:** George Q. Fei, Victoria M. Marando, Kellen L. Olszewski, Bettine E. Gibbs, Wyatt G. Jensen, Alexander M. J. Hall, Marco Jost, Suzanne Walker

**Affiliations:** Department of Microbiology, Harvard Medical School, Boston, MA, 02115, USA; Kadmon Corporation, New York, NY, 10016, USA

## Abstract

*O*-GlcNAc transferase (OGT) is an essential mammalian glycosyltransferase that installs *O*-GlcNAc onto thousands of cytosolic and nuclear proteins. Dysregulated glycosylation by OGT is associated with disease states, including cancer and neurodegeneration. However, cellular consequences of acute loss of *O*-GlcNAc remain poorly understood. Here, we employed a cell-permeable small molecule inhibitor of OGT, OSMI-4, coupled with transcriptomic and lipidomic analyses to characterize the effects of *O*-GlcNAc depletion in human cell lines. Loss of *O*-GlcNAc led to changes in expression of lipid metabolism genes and remodeling of the lipidome, with unsaturated phosphatidylcholines (PCs), saturated lysophosphatidylcholines (LPCs) and unsaturated storage lipids accumulating. We also observed increased lipid droplets. Our results suggest that polyunsaturated fatty acids (PUFAs) accumulate in the membrane under low *O*-GlcNAc conditions and that cells buffer this accumulation by removing PUFAs from membrane lipids and sequestering them in lipid droplets. We found that treatment with OSMI-4 sensitizes cells to ferroptosis inducers, likely because accumulated cell membrane PUFAs undergo peroxidation. Taken together, our results highlight lipid remodeling as an important process that mediates the downstream effects of loss of *O*-GlcNAc. It may be possible to exploit the synergy between OGT inhibitors and ferroptosis inducers for therapeutic benefit.

## Introduction

*O*-linked β-*N*-acetylglucosamine (*O*-GlcNAc) is a eukaryotic post-translational modification that consists of a GlcNAc moiety attached to a serine or threonine hydroxyl on proteins in the nucleus, cytoplasm, and mitochondria.^1–4^ The modification is catalyzed by a single enzyme, *O*-GlcNAc transferase (OGT),^5,6^ using UDP-GlcNAc as a cofactor and is also hydrolyzed by a single enzyme, *O*-GlcNAcase (OGA).^7^ Together, OGT and OGA control glycosylation of over 8000 substrates across diverse cell types and regulate a wide range of cellular processes; such as central metabolism, stress responses, and protein stability.^8,9^ Perhaps unsurprisingly, OGT is an essential protein in mammals.^10^ Transgenic mice lacking *Ogt* do not survive development, and deletion of the gene in dividing cells leads to cell death.^6,11^

One of the major hypotheses for the driving cellular function of *O*-GlcNAc is that of a nutrient sensor.^12–14^ Global *O*-GlcNAc levels are sensitive to glucose availability, and *O*-GlcNAcylation modulates activity of target substrates in a nutrient-responsive manner. Through the many target substrates of OGT, *O*-GlcNAc is thought to integrate nutrient availability into the regulation of activity for many cellular pathways, including fundamental pathways like transcription and translation. Numerous previous studies have documented individual instances of the regulation of lipid metabolism or signaling processes by *O*-GlcNAc. For example, *O*-GlcNAcylation of ATP citrate lyase (ACLY) enhances its enzymatic activity and promotes lipogenesis in cancer cells.^15^ Moreover, indirect activation of sterol regulatory element-binding protein 1 (SREBP-1) via GlcNAcylation of AMPK leads to accumulation of free fatty acids in breast cancer.^16^ Lipid signaling pathways are also modulated by *O*-GlcNAc, as in the case of the signaling lipid sphingosine-1-phosphate, whose downstream signaling is augmented by *O*-GlcNAcylation of myosin phosphatase target subunit 1 (MYPT1).^17^ The role of OGT in lipid metabolism has been further corroborated in untargeted studies. For instance, using OGT inhibitors, we previously found that perturbing *O*-GlcNAc leads to altered abundance of proteins important in sterol biosynthesis.^18^ Though several lines of evidence suggest a possible role for O-GlcNAc in regulating sterol or lipid biosynthesis, direct information on how *O*-GlcNAc levels impact the lipidome is limited. For example, it is not known if some classes of lipids are more responsive to changes in *O*-GlcNAc levels than others or if *O*-GlcNAc regulates chemical features of lipids such as fatty acid chain length or degree of unsaturation.

Here, to address this gap in our understanding, we employed the specific OGT inhibitor OSMI-4^18^ (**Fig. 1A**) to assess how human cells respond to low *O*-GlcNAc conditions. We found that highly unsaturated neutral lipids along with unsaturated phosphatidylcholines (PCs) and saturated lysophosphatidylcholines (LPCs) accumulated, as did lipid droplets. These changes can be explained if polyunsaturated fatty acyl chains accumulate in PCs under low *O*-GlcNAc conditions and are subsequently hydrolyzed and transferred to neutral lipids for sequestration in lipid droplets. Polyunsaturated lipids are prone to peroxidation by reactive oxygen species (ROS), leading to cell death by ferroptosis. Consistent with this model, we found that OSMI-4-treatment sensitizes cells to ferroptosis inducers, and this sensitivity is exacerbated by disrupting the hydrolysis of polyunsaturated fatty acids (PUFAs) from membrane lipids. We propose that sequestration of PUFAs in neutral lipids constitutes a homeostatic response that provides partial protection against ferroptosis resulting from peroxidation of accumulated membrane PUFAs. Taken together, our results highlight the interplay between *O*-GlcNAcylation and lipid metabolism and identify opportunities to exploit this interplay to enhance synergy between OGT inhibitors and ferroptosis inducers.

**Figure 1.**
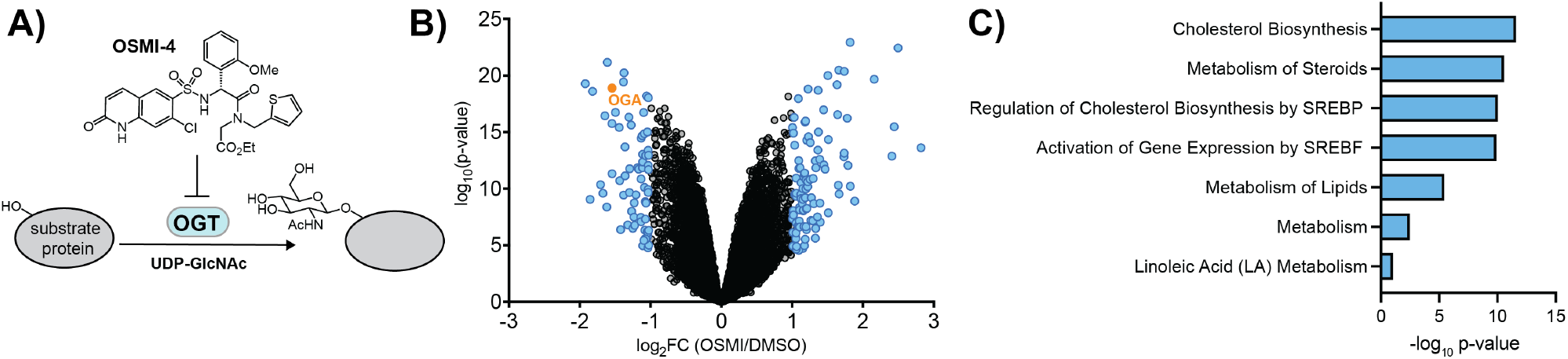
OGT inhibition changes expression of lipid metabolism and biosynthesis genes. (a) Schematic of the glycosylation reaction catalyzed by OGT showing the structure of the small molecule inhibitor OSMI-4 (previously called OSMI-4b^18^). (b) Volcano plot showing changes in transcript abundances in HCT 116 cells treated with 20 µM OSMI-4 for 24 h. Transcripts that changed at least two-fold and met our statistical cutoff (p<0.05) are represented by blue dots. OGA is highlighted in orange. (c) Top enriched Reactome terms among differentially expressed genes in OSMI-4 treated cells.

## Results

### Transcriptional responses to 24 h of OGT inhibition are primarily changes in lipid biosynthesis and metabolism

This study started with an analysis of changes to the transcriptome when OGT was inhibited for 24 h. We treated HCT 116 cells, a human colorectal carcinoma cell line, with OSMI-4 and quantified transcriptomic responses by RNA-Seq (**Fig. 1B, Fig. S1**). In total, 96 genes were upregulated and 66 genes were downregulated compared with the vehicle control (**Fig. 1B, Table S1, Table S2)**. As expected, one of the most strongly downregulated genes was OGA, which removes *O*-GlcNAc from proteins. Further analysis showed that genes involved in lipid biosynthesis, metabolism, and trafficking were highly enriched among the differentially expressed genes (**Fig. 1C, Table S3**). For example, the top two enriched Reactome pathways were “Cholesterol Biosynthesis” and “Metabolism of Steroids,” capturing the overrepresentation of genes involved in sterol synthesis and transport (*FDFT1, MVD, MVK, STARD4, DHCR7, LSS*) among the upregulated transcripts. Other top enriched Reactome pathways were “Metabolism of Lipids” and “Activation of Gene Expression by SREBF,” reflecting the overrepresentation of genes involved in the processing and trafficking of fatty acids and lipids (*SCD, FADS1, LPIN1, LDLR, ELOVL6, ELOVL3, MBOAT1*). These data suggest that OSMI-4 treatment affects lipid homeostasis across several pathways spanning biosynthesis, metabolism and trafficking, and motivated experiments to characterize changes in the lipidome following OGT inhibition.

### Cells accumulate lysophospholipids and highly unsaturated neutral storage lipids when OGT is inhibited

To characterize changes to the lipidome when OGT is inhibited, we treated HCT 116 cells with 20 µM OSMI-4 for 24 h, extracted lipids, and performed liquid chromatography-mass spectrometry-based lipidomic analysis. We identified 467 lipid species largely consisting of phospholipids, neutral lipids, and sphingolipids. When comparing OSMI-4 treated cells to DMSO controls, class-level changes in lipid abundance were observed among cholesteryl esters (CEs), triacylglycerols (TAGs), and lysophosphatidyl cholines (LPCs) (**Fig. 2A, Table S7**). We calculated an abundance metric for each lipid class by taking the geometric mean of the fold changes of all species within that class (**Table S4**). This analysis revealed that glycerophospholipid levels largely remained unchanged upon OSMI-4 treatment, whereas their corresponding lysophospholipids all displayed increases in abundance (mean fold-changes greater than 1), with LPC levels having an abundance metric of 3.5, the greatest of all lipid classes measured. TAG levels were also elevated upon OSMI-4 treatment, with an abundance metric of 1.5.

**Figure 2:**
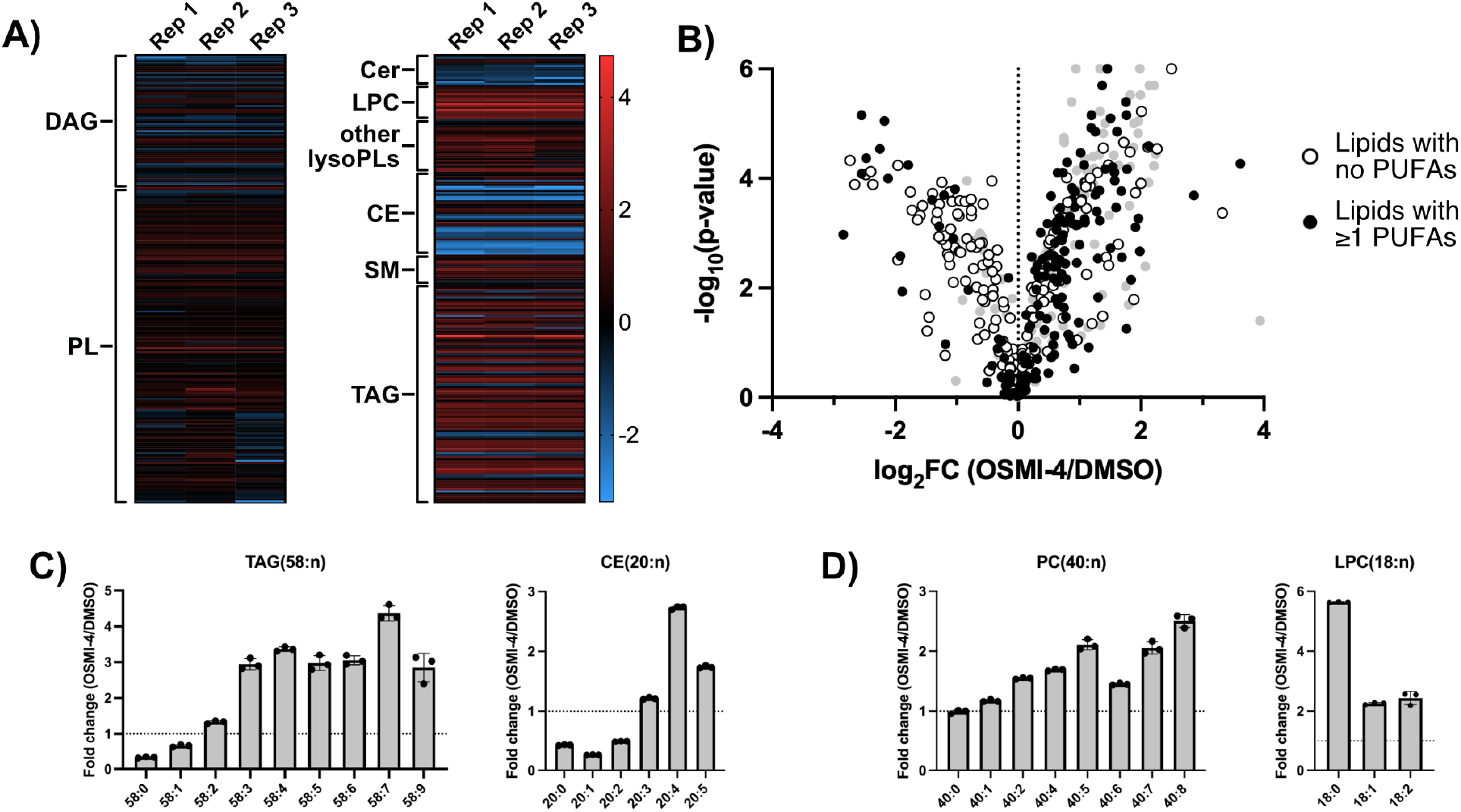
OGT inhibition leads to accumulation of unsaturated lipids. Lipidomic profile of HCT 116 cells treated with 20 µM OSMI-4 for 24 h. Lipidomic profiling was performed with n=3 biological replicates. (a) Heat map of lipid species color coded by fold-change in abundance after OSMI-4 treatment for each replicate. Fold-changes were calculated with respect to DMSO control. DAG: diacylglycerol, PL: phospholipid (including PC: phospatidylcholine, PE: phosphatidylethanolamine, PG: phosphatidylglycerol, PI: phosphatidylinositol and PS: phosphatidylserine), Cer: ceramides, LPC: lysophosphidylcholine, other lysoPLs: other phospholipids (including LPE: lysophosphatidylethanolamine, LPI: lysophosphatidylinositol, LPS: lysophosphatidylserine), CE: cholesteryl ester, SM: sphingomyelin, TAG: triacylglycerol. (b) Volcano plot with each dot representing a lipid species from lipidomic profiling. Lipid species guaranteed to have no PUFAs are represented by white dots, whereas species guaranteed to have at least one PUFA are represented by black dots (See methods). Species that may or may not have any PUFAs are represented by grey dots. (c)-(d) Graphs showing fold-changes in selected lipid classes that highlight trends in saturation state.

Lipid species within a class were further grouped by number of acyl carbons, which revealed a pattern in which lipid species containing only saturated or monounsaturated fatty acids tended to be depleted, while highly unsaturated species tended to be enriched (**Fig. 2B**). For neutral lipids (DAGs, TAGs, and CEs), among species with the same number of carbons, the fold-change in abundance upon OSMI-4 treatment increased with increasing degree of unsaturation. Taking TAG(58:n) lipids as an example, TAG(58:0) and TAG(58:1) decreased significantly compared to the DMSO treatment, while TAG species with more unsaturation (58:3 and beyond) increased significantly (**Fig. 2C**). Similarly for CE(20:n) lipids, species with low unsaturation such as CE(20:0), CE(20:1), and CE(20:2) decreased in abundance while CE(20:4) and CE(20:5) increased in abundance upon OSMI-4 treatment compared to DMSO (**Fig. 2C** and **Fig. S2**). DAGs displayed the same pattern of unsaturation, but the changes were not as large, perhaps reflecting their role as intermediates between phospholipids and TAGs. We also calculated a score based on the average degree of unsaturation per acyl chain weighted by the fold-change of each individual species in that class (**Table S5**). TAGs, CEs, and DAGs had the largest increase in their unsaturation scores. These data are consistent with a shift in the synthesis of neutral lipids towards species containing PUFAs.

With the exception of PCs, most phospholipids did not show a pattern in their unsaturation levels upon OGT inhibition. For PCs, however, species with more than one degree of unsaturation tended to increase in abundance (**Fig. 2D** and **Fig. S2**). For example, among PC(40:n) lipids, PC(40:2) through PC(40:8) all showed significant increases in abundance. These observations were also corroborated by the unsaturation score analysis. Notably, LPCs, which are related to PCs via removal of the *sn*-2 acyl chain, tended to show the opposite pattern of unsaturation. Even though all detected LPC species were more abundant in the OSMI-4 treated condition, saturated LPCs generally increased more than unsaturated species. For example, LPC(18:0) increased almost 6-fold upon OSMI-4 treatment, while LPC(18:1) and LPC(18:2) only increased by 2-fold (**Fig. 2D**). In our unsaturation score analysis, LPCs were the only lipid species to trend towards a lower degree of unsaturation. Overall, these data provide evidence that OGT impacts the levels of several lipid classes and may regulate the PUFA content of cellular lipids.

The HCT 116 cell line used for the lipidomic experiments was also engineered to express a degron-tagged OGT variant (dTAG-OGT) from the endogenous *OGT* locus, enabling the rapid depletion of OGT upon treatment with a degrader compound (dTAG-v1) as an orthogonal means of perturbing *O*-GlcNAc levels. Lipidomic analysis using the degrader compound showed that OGT protein depletion recapitulated lipid abundance changes across lipid classes, LPCs again showing the greatest increase in abundance (**Fig. S3, Table S6, Table S7**). Although the changes for the degrader compound were not as large as for the inhibitor, likely because degradation leaves a pool of active OGT,^19^ the results with the degrader compound serve as orthogonal validation that reducing *O*-GlcNAc remodels the lipidome.

### OGT inhibition induces lipid droplet accumulation independent of fatty acid synthesis

Neutral lipids are primarily found in lipid droplets (LDs), storage organelles comprising a phospholipid monolayer surrounding a hydrophobic core predominantly composed of TAGs and CEs.^20^ We hypothesized that the accumulation of highly unsaturated TAGs upon inhibition of OGT would manifest as an increase in the number of LDs present in the cell. To visualize LDs, we stained cells using BODIPY 493/503, a lipophilic dye that preferentially partitions into LDs, and imaged the cells by confocal microscopy. The number of BODIPY-stained puncta increased in OSMI-4-treated cells compared to the DMSO control, in line with our lipidomics data (**Fig. 3A and 3B**). An enantiomer of OSMI-4 that binds more weakly to OGT, OSMI-4ent,^18^ failed to increase LD numbers per cell compared to DMSO, confirming that the accumulation of LDs is due to on-target effects of OSMI-4 rather than to non-specific effects of the relatively hydrophobic OSMI scaffold (**Fig. S4A**). OSMI-4 also caused LD accumulation in U2OS cells, an osteosarcoma cell line (**Fig. S4B**), indicating that the effect is not limited to HCT 116 cells. To verify that the LD phenotype is indeed a result of TAG accumulation, we used T863, a small molecule inhibitor of diacylglycerol acyltransferase 1 (DGAT1), which is primarily responsible for producing TAGs from DAGs (**Fig. 3C**).^21^ Treatment of cells with T863 lowered the baseline number of BODIPY-stained puncta and also abolished the OSMI-4-dependent increase in puncta compared to the DMSO control (**Fig. 3D**), confirming that the accumulation of LDs induced by OGT inhibition depends on TAG synthesis.

**Figure 3:**
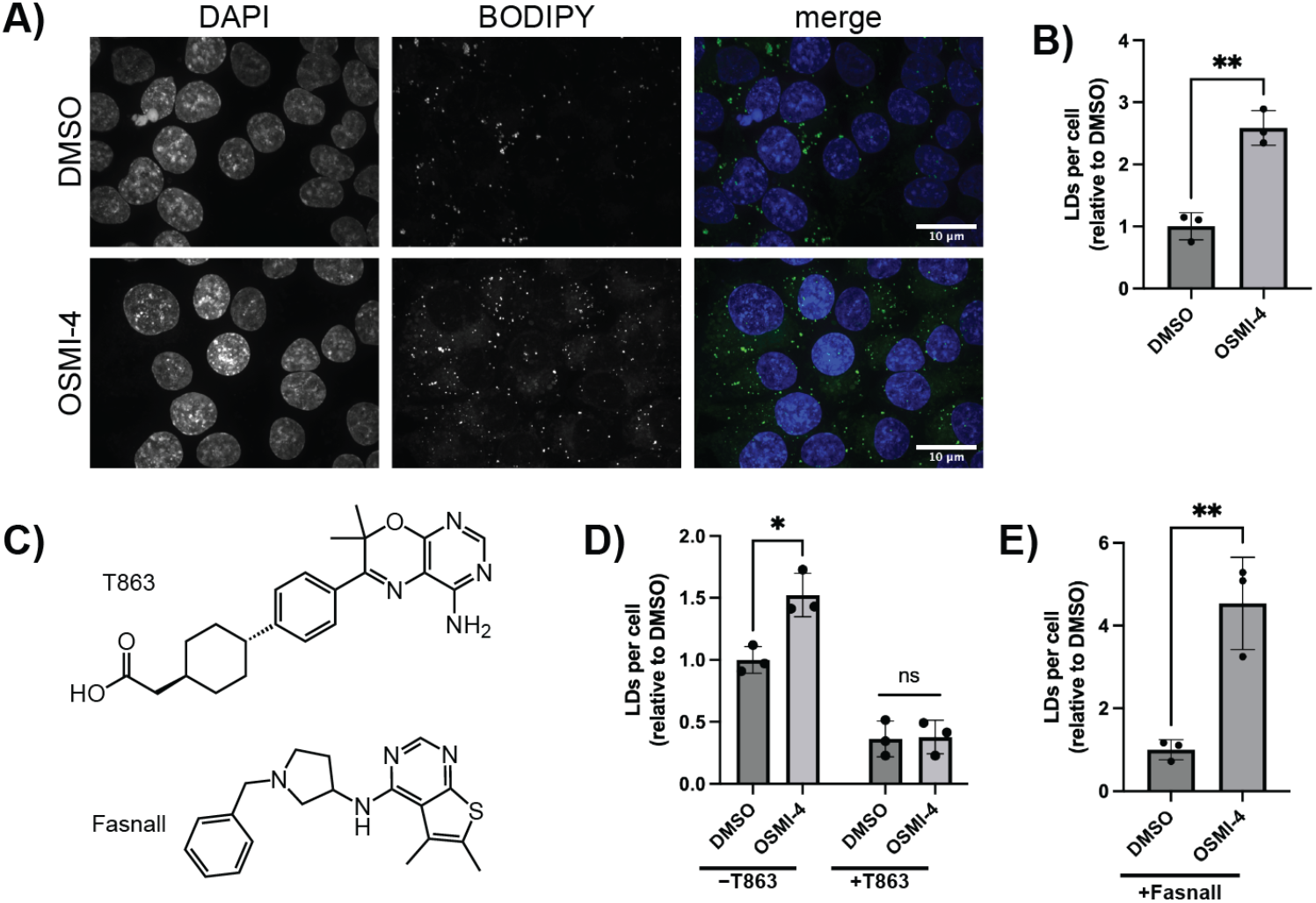
OGT inhibition induces lipid droplet formation. Confocal microscopy analysis of LD abundance in HCT 116 cells. Analysis was performed with *n*=3 biological replicates. Quantification was performed on maximum intensity projections. (a) Confocal microscopy images of HCT 116 treated with DMSO or 20 μM OSMI-4 for 24 h. (b) Quantification of LDs per cell in HCT 116 cells treated with 20 μM OSMI-4. Data was normalized relative to DMSO. (c) Chemical structures of small molecule inhibitors used. T863 inhibits DGAT1 (diacylglycerol O-acetyltransferase 1). Fasnall inhibits FASN (fatty acid synthase). (d) Quantification of LDs in HCT 116 cells co-treated with 20 μM OSMI-4 and 2 μM T863 for 24 h. (e) Quantification of LDs in HCT 116 co-treated with 10 μM fasnall and 20 μM OSMI-4 for 24 h.

We considered the possibility that *de novo* synthesis provides fatty acids for TAG accumulation. This seemed unlikely because highly unsaturated lipids were found in the TAGs, and human cells cannot synthesize PUFAs *de novo*, although they contain fatty acid desaturases that further increase unsaturation of PUFAs obtained from dietary sources.^22^ Nevertheless, to test the role of *de novo* synthesis in LD accumulation, we inhibited fatty acid synthase, the key enzyme in the pathway, with fasnall (**Fig. 3C**).^23^ In the presence of fasnall, OGT inhibition still resulted in an increase in the number of LDs (**Fig. 3E**), implying that the lipids that accumulate during low *O*-GlcNAc conditions are derived from extracellular sources. Treatment of HCT 116 cells grown in media supplemented with delipidated serum resulted in a dose-dependent growth defect, indicating on-target activity of fasnall in our system (**Fig. S4D**).

### OGT inhibition sensitizes cells to ferroptosis

Lipids containing PUFAs are prone to peroxidation by reactive oxygen species (ROS), with runaway lipid peroxidation leading to a form of cell death known as ferroptosis (**Fig. 4A**).^24^ Accumulation of membrane PUFAs is known to sensitize cells to pro-ferroptotic redox stress, and sequestration of PUFAs into lipid droplets has been proposed to be a homeostatic mechanism to buffer against ferroptosis.^25,26^ Based on our lipidomics and lipid droplet microscopy data, we hypothesized that OSMI-4 treatment causes ferroptotic stress by an accumulation of PUFAs. We expected that if this were the case, HCT 116 cells treated with OSMI-4 would be more sensitive to pharmacologic induction of ferroptosis. To test this, we dosed cells with ML210, a covalent GPX4 inhibitor used widely as a ferroptosis inducer^27^(**Fig. 4A/B**). We found that treatment with OSMI-4 sensitized cells to ML210 compared to DMSO (**Fig. 4C**). This sensitization was abolished by co-treatment with ferrostatin-1, a radical scavenger that inhibits ferroptosis by preventing lipid peroxidation^28^ (**Fig. 4C**), demonstrating that the synergy between OSMI-4 and ML210 is dependent on ferroptosis.

**Figure 4:**
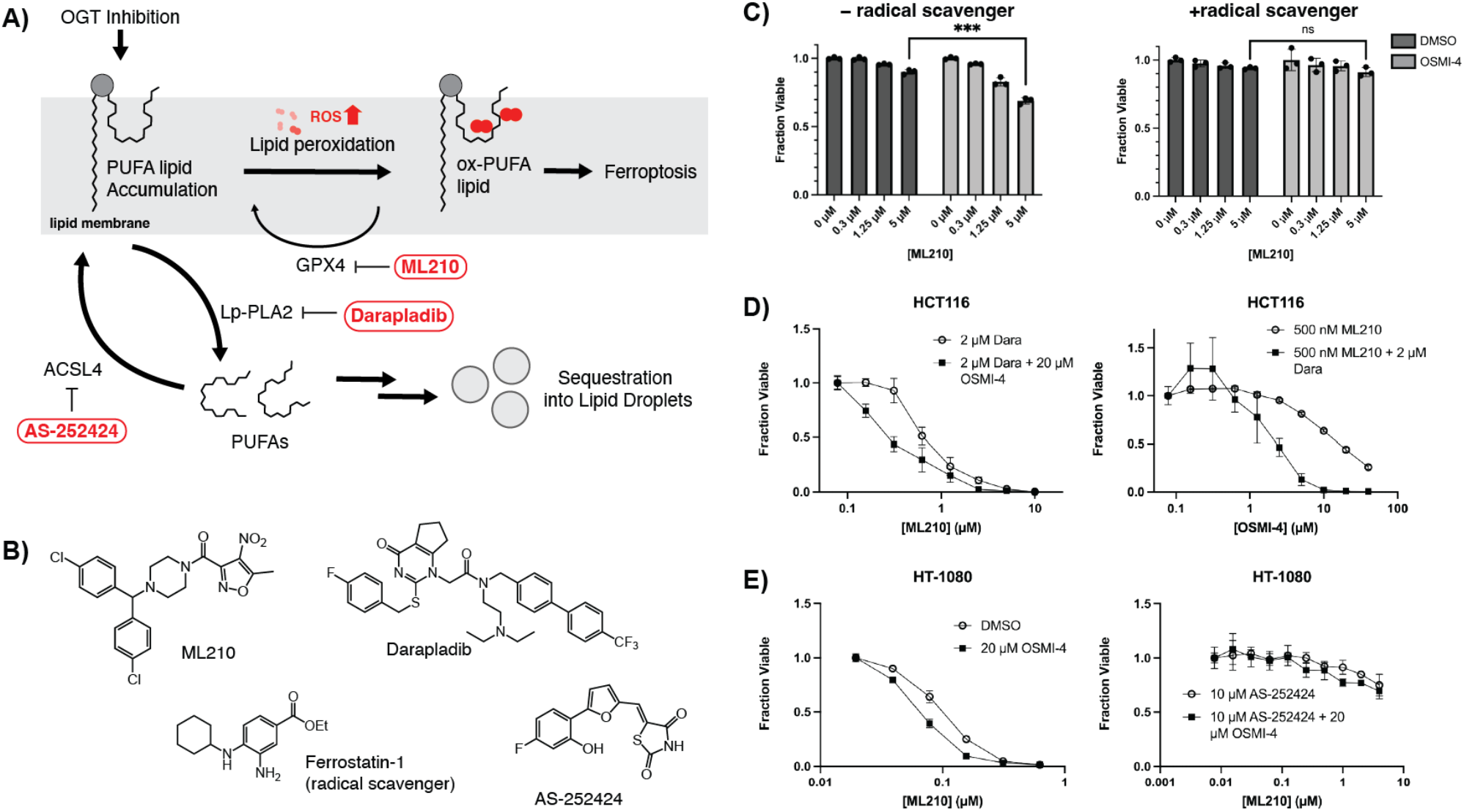
OGT inhibition sensitizes cells to ferroptosis. (a) Working model showing that cells treated with OSMI-4 accumulate membrane PUFAs, leading to a pro-ferroptotic state. Channeling PUFAs from the cell membrane into lipid droplets serves as a buffering mechanism against ferroptosis. Small molecule inhibitors described in the text are highlighted. ML210 targets GPX4 (glutathione peroxidase 4), an enzyme that reduces lipid peroxidation. Darapladib inhibits Lp-PLA2 (lipoprotein-associated phospholipase 2), which hydrolyzes membrane-resident PUFAs. AS-252424 inhibits ACLS4 (acyl-CoA synthetase long chain family member 4), which converts PUFAs into active PUFA-CoA intermediates to enter metabolic or remodeling pathways. (b) Chemical structures of small molecules used to perturb ferroptosis-related pathways. (c) Dose-response of HCT 116 cells to ML210 with or without 20 μM OSMI-4 and 15 μM Ferrostatin-1. Data points indicate 3 biological replicates; error bars indicate mean +/-S.D. (d) Dose-response curve of HCT 116 to ML210 in the presence of 2 μM darapladib, with or without 20 μM OSMI-4. *n*=3 biological replicates; error bars represent mean +/-S.D. (e) Dose-response curve of HT-1080 cells to ML210 in the presence of 10 μM AS-252424, with or without 20 μM OSMI-4. *n*=3 biological replicates; error bars represent mean +/-S.D.

The phospholipase Lp-PLA2 hydrolyzes PUFAs from membrane lipids and protects cells against ferroptosis.^29^ We hypothesized that if OSMI-4 treated cells were in a ferroptosis-prone state, they would be reliant on Lp-LPA2 activity to maintain viability. We used the Lp-PLA2 inhibitor darapladib to test whether disrupting removal of PUFAs from the membrane would enhance the synergy between ML210 and OSMI-4 (**Fig. 4A/B**). Darapladib treatment sensitized HCT 116 cells to ML210 alone (**Fig. 4D**), as expected. In the presence of darapladib, ML210 inhibited growth of HCT 116 cells with a submicromolar IC_50_, whereas without darapladib, ML210 did not fully inhibit cell growth even at 20 µM, the highest concentration we were able to test (**Fig. 4D** and **Fig. S5**). Addition of OSMI-4 further sensitized cells to the combined effect of ML210 and darapladib (**Fig. 4D**). We concluded that OSMI-4 synergizes with ML210 and that sequestration of membrane PUFAs protects cells. Consistent with this, treatment of HCT 116 cells with ML210 increased the number of LDs per cell, and co-treatment with both ML210 and OSMI-4 resulted in a larger number of LDs compared to just ML210 or OSMI-4 alone (**Fig. S6**).

We next wanted to test whether preventing the incorporation of PUFAs into membrane lipids would reduce the sensitization to ferroptosis by OSMI-4. Because HCT 116 cells are not very sensitive to ML210 treatment, we switched to HT-1080 cells, a human fibrosarcoma cell line naturally possessing submicromolar sensitivity to many GPX4 inhibitors.^30^ We confirmed that in HT-1080 cells, OSMI-4 treatment resulted in loss of *O*-GlcNAc (**Fig. S7**), led to LD accumulation (**Fig. S8**), and sensitized cells to ML210 (**Fig. 4E**). In contrast, OGA inhibition did not sensitize cells to ML210 (**Fig. S9**). To prevent PUFA incorporation, we used an inhibitor of ACSL4, the acyl-CoA ligase specific for arachidonic acid, a PUFA known to be vulnerable to peroxidation (**Fig. 4A**).^31^ The ACSL4 inhibitor AS-252424 has been shown to be protective against ferroptosis inducers by reducing membrane PUFA content and overall peroxidation of membrane lipids (**Fig. 4B**). When treated with AS-252424, the HT-1080 cells were insensitive to ML210 and did not display an OSMI-4-dependent sensitization to ML210 (**Fig. 4E**), implying that the synergy between ML210 and OSMI-4 depends specifically on membrane PUFAs.

## Discussion

We have shown here that *O*-GlcNAc plays a critical role in regulating lipid metabolism transcriptionally and modulating lipidome composition. Cells with low *O*-GlcNAc accumulate saturated LPCs, unsaturated PCs, and unsaturated neutral lipids. Neutral lipids, in particular triglycerides, accumulate in lipid droplets in a DGAT1-dependent manner. LPCs and PCs are related to each other by the Lands cycle, wherein fatty acids are hydrolyzed from and esterified onto membrane phospholipids, allowing cells to modify the fatty acid composition of membranes.^32^ Our data are consistent with a model in which PUFAs are hydrolyzed from PCs in the membrane, leaving behind saturated LPCs; the liberated PUFAs are then sequestered in neutral lipids stored in LDs. These lipidome-level changes may reflect a homeostatic program that ameliorates ferroptotic stress by removing peroxidizable PUFAs from membranes. Consistent with this, we found that sensitivity to a ferroptosis inducer increased upon *O*-GlcNAc inhibition, particularly when removal of the highly unsaturated lipid arachidonic acid from membrane lipids was prevented. Conversely, preventing incorporation of arachidonic acid into membrane lipids protected cells from OSMI-4-induced sensitivity to ferroptosis.

Previous work has demonstrated that OGT regulates *de novo* fatty acid synthesis by glycosylating a specific residue on ACLY.^15^ Preventing *O*-GlcNAc modification at this site reduced fatty acid and lipid quantities across multiple lipid classes. In contrast, we found that inhibiting OGT led to a specific remodeling of lipids involving PUFA-containing species. This remodeling is not mediated through *de novo* fatty acid synthesis, indicating that regulation of PUFA lipids is a distinct and previously unappreciated function of *O*-GlcNAc. Previous work has also shown that in visceral fat adipocytes, loss of *O*-GlcNAc destabilizes the key LD factor perilipin 1 (PLIN1), thereby promoting lipolysis and causing shrinkage of LDs.^33^ We found, instead, that loss of *O*-GlcNAc leads to an increase in LDs across a panel of mitotically active cultured cell lines. This discrepancy can be explained by the adipocyte-specific expression of PLIN1^34^, with the influence of *O*-GlcNAc on LDs likely mediated through different factors in non-adipocyte cell types. For example, consistent with our observations, another group found that in HT-29 cells, a human colorectal cancer cell line, knockout of *OGT* or replacement with a catalytically inactive variant results in an increase in LD content.^35^ They showed that this occurs in part via induction of PLIN2, a homolog of PLIN1. Together with our data, these studies underscore how OGTs role in regulating LDs changes depending on the cellular context. Rapidly dividing cancer cell lines may share related pathways for regulation of their LDs, whereas adipocytes, due to their specialized role in systemic lipid metabolism, employ distinct mechanisms for managing LD content. Further studies are needed to determine whether OGT regulates LD dynamics in other cell types with specialized lipid-processing roles, such as hepatocytes, cardiomyocytes, and macrophages. Additionally, the impact of O-GlcNAc on levels of free cholesterol and cholesterol precursors should be examined in future studies because they were not captured in the lipidomics detection platform used here.

Based on the studies described here, we present a working model in which OSMI-4-induced PUFA dysregulation leads to sensitivity to ferroptosis inducers. Lipidomic analysis showed that PCs tend to become more unsaturated upon OSMI-4 treatment. Indeed, the PCs that displayed the greatest fold-changes were the most unsaturated, like PC(40:8). This species likely represents a double-PUFA lipid, which are known to be drivers of ferroptosis.^36^ In our transcriptomic analysis, OSMI-4 treatment also resulted in an increase in *FADS1* expression and a decrease in *MBOAT1* expression. Both of these genes are involved in regulating the abundance of PUFA-containing phospholipids, and an increase in FADS1 and decrease in MBOAT1 would predict an accumulation of PUFA-containing phospholipids, consistent with our lipidomics data.^37,38^

While this data points to altered PUFA metabolism as the primary driver of ferroptotic sensitivity under low *O*-GlcNAc conditions, other mechanisms may contribute to this sensitivity. The lipid peroxidation chain reaction that characterizes ferroptosis requires labile iron (II) and ROS.^26^ A previous study implicated increased labile iron (II) liberated through ferritinophagy and mitophagy.^39^ Moreover, *O*-GlcNAc is known to regulate the redox state of the cell, and ROS production may exacerbate ferroptotic sensitivity when OGT is inhibited. Further study is required to elucidate the requirement for ROS production to initiate ferroptosis in the context of OGT inhibition.

Ferroptosis has attracted substantial attention since its discovery owing to its potential as a targetable anti-tumor pathway.^24,40^ Though certain cancer types are especially susceptible to ferroptosis stress, many types exhibit little susceptibility.^41^ Inhibition of OGT offers a potential avenue for sensitizing cells to ferroptosis inducer treatment or even widening the therapeutic window to prevent induction of ferroptosis in healthy tissue. Additionally, disruption of *O*-GlcNAc cycling is being explored as a therapeutic strategy for cancers as well.^42^ Our data show that lipid remodeling is an important process that mediates downstream effects of loss of *O*-GlcNAc and that disruption of this remodeling, specifically through inhibition of Lp-PLA2 using darapladib, can augment the synergy between OGT inhibitors and ferroptosis inducers.

## Acknowledgements

We thank the Core for Imaging Technology and Education at the Harvard Medical School for microscopy instrumentation and expertise. We thank Dr. Magdalena Bach and Dr. Nika Danial and for their counsel regarding ferroptosis. Research reported in this work was supported by the National Institute of Health under grant number R01GM094263 and R01AI148752 (to S. W.), NIH grant R01GM094263-11S1 (to B. E. G.) and NIH grant DP2GM154152 (to M. J.). This work was also supported by the Helen Hay Whitney Foundation fellowship (to V. M. M). A total of 80% of this project was financed with federal funds. A total of 20% of was financed with non-governmental funds.

## Author Contributions

G. Q. F. and S. W. designed research with input from K.O. and V.M.M. G. Q. F., V. M. M., K. O., W. G. J. and A. M. J. H. conducted research. G. Q. F., V. M. M., and K. O. analyzed data. B. E. G. contributed new reagents. M. J. provided support and guidance. G. Q. F., V. M. M., and S. W. wrote the paper with input from all authors.

## Disclaimer

The content is solely the responsibility of the authors and does not necessarily represent the official views of the National Institute of Health.

## Data Availability

RNAseq data is deposited in the GEO under accession GSE352806. https://www.ncbi.nlm.nih.gov/geo/query/acc.cgi?acc=GSE352806

## Supporting Information

Additional experimental details, materials, and methods as well as supplementary figures and tables containing experimental results.

## TOC Graphic

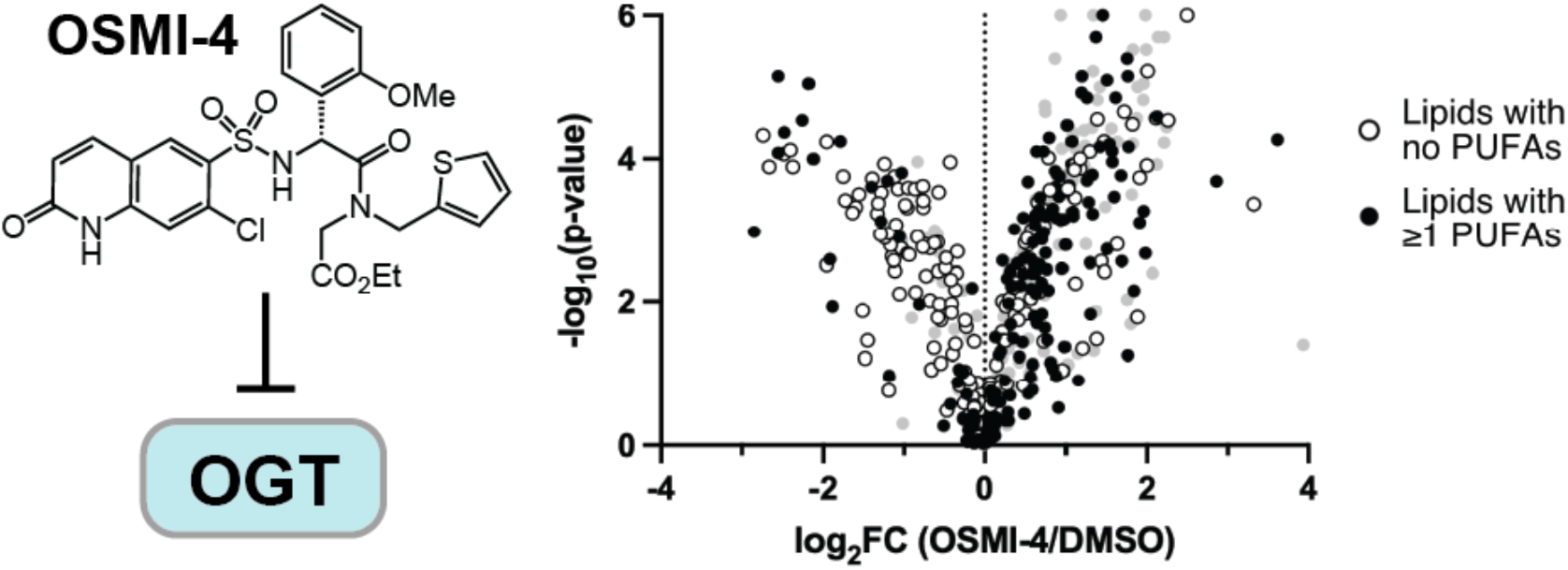

