## Supporting information for "Lipid Changes due to Low O-GlcNAc and the Downstream Consequences"

### Supplemental Procedures

**Cell Line Maintenance**

HCT 116 WT and HCT 116 dTAG-OGT cells were cultured in RPMI 1640 media with GlutaMAX$^{TM}$ supplement (Thermo Fisher 72400120). U2OS cells were cultured in McCoy's 5A (modified) media (Thermo Fisher 16600082), and HT-1080 cells were cultured in Dulbecco's modified Eagle medium (DMEM), low glucose, with GlutaMAX$^{TM}$ and pyruvate supplement (Thermo Fisher 10567-022). All media was supplemented with 10% fetal bovine serum (Thermo Fisher A5256701). All cells were cultured in incubators set to 37°C with 5% $CO_2$.

For compound treatment experiments, cells were plated in 6-well plates at a density of 500,000 cells per well. After 24 h, media was removed, and 2 mL of fresh media was added to each well. Cells were treated with DMSO (Millipore Sigma D8418) or compound for the indicated amount of time. Compounds were obtained from the following sources: T863 (MedChem Express HY-32219), fasnall (MedChem Express HY-121250), ML210 (MedChem Express HY-100003), ferrostatin-1 (MedChem Express HY-100579), darapladib (MedChem Express HY-10521), AS-252424 (Cayman Chemical Company 10009052), dTAG-V1 (Tocris 6914), OSMI-4 was synthesized in-house following literature procedures.[1]

**RNA-Seq**

HCT 116 cells were seeded in a 24-well tissue culture treated plate at 100K cells per well. 5 replicates of DMSO treatment and 5 replicated of 20 μM OSMI-4 treatment were prepared. Following a 24 h incubation, cells were harvested by treatment with TrypLE Express (ThermoFischer 12605036), pelleted by centrifugation at 500 x*g* for 5 min and the resultant pellets were resuspended in 50 μL of Zymo DNA/RNA Shield™ (Zymo Research R1100-50). Lysates were submitted to Plasmidsaurus for RNA-Seq analysis using Illumina sequencing and a 3' end counting approach. Hits were defined as having a $\log_2FC > |1|$, p-value $< 0.05$ and logCPM $> 1$. Gene Set Enrichment Analysis run using GSEApy[2] and the Enrichr library[3] "Reactome_Pathways_2024".

**Cell Line Construction**

*Sp*Cas9 and sgRNA plasmid

Oligos targeting exon 22 proximal to the 3'UTR of OGT were synthesized and ordered from Integrated DNA Technologies. Digestion of the plasmid, oligo annealing, and ligation were completed following the protocol from Feng Zhang's lab[4] for the human codon-optimized *Sp*Cas9 and sgRNA expressing PX459 V2.0 plasmid (Addgene #62988, https://www.addgene.org/crispr/zhang/). 2 μL of the ligation reaction was directly transformed into NEB Stable competent *E. coli* following the manufacturers protocol (New England Biolabs, cat. #C3040H). Following mini-prep, the plasmid was sequenced and subsequently midi-prepped prior to transient transfection.

Oligo #1 5'—CACCGCCTGAGCCTCAACCTTCTGG—3'
Oligo #2 3'—CGGACTCGGAGTTGGAAGACCCAAA—5'

Donor plasmid
The empty backbone pFETCH Donor plasmid (Addgene #63934) was used to generate the OGT dTAG[5] knock-in largely following the protocol from the Mendenhall and Myers lab groups (https://media.addgene.org/data/plasmids/63/63934/63934-attachment_HGifbVXT3xET.pdf). The NeoR cassette was swapped for sfGFP. Briefly, homology arms were ordered as gBlocks from IDT and PCR amplified using Q5 High-Fidelity 2X Master Mix (New England Biolabs, cat. #M0492S). The amplicons were cleaned up with Qiagen PCR purification and used in a Gibson Assembly reaction with the digested pFETCH vector and a PCR amplified sfGFP fragment to generate the OGT-targeting plasmid**.**

Transfection
Wild-type HCT 116 cells were plated in a 6-well dish at a density of 300,000 cells/2.5 mL and incubated for 18-24 hours prior to transfection. Using a co-expression method 2.5 μg of DNA was transiently transfected into HCT 116 cells using the TransIT XT-2 reagent (Mirus, SKU: MIR 6004) with a 1:1 ratio of Cas9/sgRNA plasmid to Donor plasmid.

Cells were then sorted from the top 0.5% of sfGFP+ into a 96-well plate in 100 μL of media where the first well (A1) contained 100 cells and each subsequent well had ~1 cell/well. Only in wells where single colonies were formed was media aspirated and 50 μL of TrypLE used to detach cells from the well. Cells were moved from the 96-well plate to a 12-well plate and finally a 6-well plate where they were split and gDNA was extracted for orthogonal PCR genotyping.

PCR genotyping
Oligo Pair, Junction 1
Fwd: 5' CTGAACTCAGTTCAGGTACAAG 3'
Rev: 3 ' GTGGAGCTTCTAAAACTGGAAG 5'
Oligo Pair, Junction 2
Fwd: 5' GGAACTAGAGCGGCTCTATC 3'
Rev: 3' CAGTACCTTGTTGCAGATG 5'
Positive clones were then expanded and subsequent immunoblot analysis was completed.

**Lipidomics Sample Preparation and LC-MS**
After compound treatment, media was aspirated from the 6-well plates, and the cells were washed with 1 mL of room temperature PBS. After removal of the PBS, plates were placed on ice, and 1 mL of ice-cold PBS was added to each well. Cells were detached by scraping and pelleted in a centrifuge at 1000×*g* for 1 min at 4°C. After discarding the supernatant, cells were resuspended in 250 μL cold isopropanol. The tubes were placed on wet ice and lysed by sonication for 5 min. The homogenates were centrifuged at 21000 x*g* for 20 min at 4°C, and the supernatants were transferred to pre-chilled autosampler vials.

Lipidomics analysis by LC–MS was performed using an analytical method based on a method previously described.[6] The extracts were transferred to amber glass sample vials in an autosampler maintained at 4 °C. Separation was performed on a Thermo Scientific UltiMate 3000 HPLC system using an Acquity UPLC HSS T3 column (2.1 × 100 mm, 1.8 μm particle size). The HPLC parameters are as follows: Mobile phase A (acetonitrile:water (60:40, v/v) with 10 mM ammonium acetate and 0.1% acetic acid), Mobile phase B (isopropanol:acetonitrile:water (85:10:5, v/v/v) with 10 mM ammonium acetate and 0.1% acetic acid), Gradient (0 min, 40% B; 1.5 min, 40% B; 12 min, 100% B; 15 min, 100% B; 16 min, 40% B; 17 min, 40% B), Injection volume (10 µl), Column temperature (55 °C), and Flow rate (400 µl/min).

Samples were analyzed on an Exactive orbitrap mass spectrometer in both positive and negative ionization mode with a heated electrospray ion source. The instrument parameters are as follows: sheath gas flow rate 30 (arbitrary units), aux gas flow rate 10 (arbitrary units), sweep gas flow rate 3 (arbitrary units), spray voltage 4 kV, capillary temperature 120 °C, heater temperature 500 °C, capillary voltage 65 V, tube lens voltage 100 V. The scan range was set to 200–1,000 *m*/*z*, with a maximum inject time of 100 ms, resolution of 100,000 at 1 Hz, and AGC (automatic gain control) target $1 \times 10^6$. The data were analyzed using the MAVEN software suite 57 with signal intensity determined as the Peak Area (Top). Depending on the lipid analyte, we measured either the molecular ion or the ammonium adduct ($[M+NH4]^+$).

**Lipidomics Analysis**
All lipid species were grouped into one of the following classes: ceramides (Cer), sphingomyelins (SM), cholesterol esters (CE), diacylglycerols (DAG), triacylglycerols (TAG), phosphatidylcholines (PC), phosphatidylethanolamines (PE), phosphatidylserines (PS), phosphatidylinositols (PI), phoshatidylglycerols (PG), lysophosphatidylcholines (LPC), lysophosphatidylethanolamines (LPE), lysophosphatidylserines (LPS), and lysophosphatidylinositols (LPI). Lipid classes for which less than 5 species were detected were excluded from this analysis. The overall fold change for each lipid class was calculated as the geometric mean of the fold-changes for all species in that class.

To calculate the change in saturation, lipid species within a class were grouped by number of carbons. The change in saturation due to inhibitor treatment within a group of lipids with the same carbon number was calculated using **Equation 1**,

$$\Delta_{sat} = \frac{\sum_{i \in \text{Group}} DoU_i \times FC_i}{\sum_{i \in \text{Group}} FC_i} - \frac{\sum_{i \in \text{Group}} DoU_i}{n_{\text{Group}}} \quad \textbf{(1)}$$

where $DoU_i$ is the degree of unsaturation of the *i*-th lipid species in the group, $FC_i$ is the fold-change in abundance of the *i*-th lipid species in the group after inhibitor treatment, and $n_{\text{Group}}$ is the total number of lipid species in the group. The $\Delta_{sat}$ was calculated for each group within a class, and these saturation scores were averaged across all groups within a class to generate the overall change in saturation for that class. The saturation score for a class was then normalized to the number of acyl chains of that class.

Lipids species that only had zero or one degree of unsaturation were labeled as "guaranteed to have no PUFAs," while, by the pigeonhole principle, lipid species of $n_{acyl}$ acyl chains that had at least $n_{acyl}$+1 degrees of unsaturation were labeled as "guaranteed to have at least one PUFA." Lipid species with more than one but less than $n_{acyl}$+1 degrees of unsaturation could not be assigned to either label.

**Immunoblot Analysis**
Cells were cultured in 6-well plates. After compound treatment, cells were washed with ice-cold PBS (Thermo Fisher 14190250) and detached by scraping. Cells were pelleted by centrifugation at 200×*g* for 5 min at 4°C and was frozen overnight at −80°C in a freezer and resuspended the following day in 100 μL RIPA buffer supplemented with 1X cOmplete EDTA-free protease inhibitor cocktail (Millipore Sigma Cat. No. 11873580001). The resulting lysate was spun down at 21,000×*g* for 10 min at 4°C, and 100 μL of the supernatant was mixed with 100 μL 2X Laemmli and incubated at 95°C for 5 min. Proteins were separated by electrophoresis in a 4-20% Criterion™ TGX™ precast midi protein gel (Bio-Rad Cat. No. 567109) at 170 V for 55 min. Proteins were transferred to a nitrocellulose membrane (Bio-Rad Cat. No. 1620168) using the Bio-Rad Trans-Blot® Turbo™ transfer system. The membrane was blocked at room temperature on a platform rocker for 1 h using SuperBlock™ T20 (TBS) blocking buffer (Thermo Fisher Cat. No. 37536). The membrane was incubated with the primary antibody (O-GlcNAc (RL2) Abcam ab2739; OGT Cell Signaling Technologies 24083S; OGA (MGEA5) Sigma Aldrich HPA036141; β-actin Cell Signaling Technologies 3700S) in SuperBlock™ overnight at 4°C. The membrane was washed three times in TBST and incubated in a solution of the HRP-linked secondary antibody in SuperBlock™ at room temperature for 1 h. The membrane was washed three times with TBST. The blot was imaged using Pierce™ ECL western blotting substrate (Thermo Fisher Cat. No. 32106) on a BioRad Chemidoc imager. Image quantification was carried out in BioRad Image Lab 6.1.

**Lipid Droplet Microscopy**
For microscopy experiments, cells were plated on 6-well plates with No. 1.5 glass coverslips (Mattek Cat. No. P06G-1.5-20-F). Compound treatment was the same as other experiments. After compound treatment, 1 mL of media was removed from each well, and 1 mL of 8% paraformaldehyde (Thermo Fisher Cat. No. PI28908) was added to each well. The plate was incubated at room temperature for 15 min, after which each well was aspirated and washed with 1 mL of PBS. 1 mL of 1 μg/mL DAPI was added to each well, and the plate was incubated at room temperature in the dark for 1 h, after which each well was aspirated and washed with 1 mL of PBS. 1 mL of 1 μM BODIPY 493/503 (MedChemExpress Cat. No. HY-W090090) in PBS was added to each well, and the plate was incubated in the dark at 37°C for 15 min, after which each well was aspirated and washed once with 1 mL of PBS. Stains samples were stored in 1 mL of PBS at 4°C until imaging.

Microscopy was performed on a Nikon Ti motorized inverted microscope with a Yokogawa CSU-X1 spinning disk confocal, a Hamamatsu ORCA-Fusion BT sCMOS camera, and a Nikon LUN-F XL solid state laser combiner. Cells were imaged on the 60X objective. No binning or gain was applied to the imaging process. Lipid droplets were identified using the threshold function in ImageJ and quantified using the analyze particles function in ImageJ. The same threshold was applied across all samples of an experiment.

**Dose Response Curves**
Cells were plated on white-wall, clear bottom 96-well plates (Corning Cat. No. 353377) at a density of 10,000 cells per well and were left to adhere to the plate overnight. The next day, the media was aspirated off each well, and fresh media containing the appropriate concentration of either DMSO or compound was added to each well. The cells were cultured for 48 or 72 h, after

which cell viability was assayed by CellTiter-Glo$^{R}$ (Promega Cat. No. G7570). Luminescence was measured using a ProMega GloMax$^{R}$ Discover microplate reader. Luminescence signals for a given dose series were normalized to the signal from the lowest concentration treatment. For dose response experiments carried out in the presence of another compound, luminescence signals were normalized to the signal from the lowest concentration treatment of the titrated compound in the presence of the secondary compound.

### Supplemental Figures

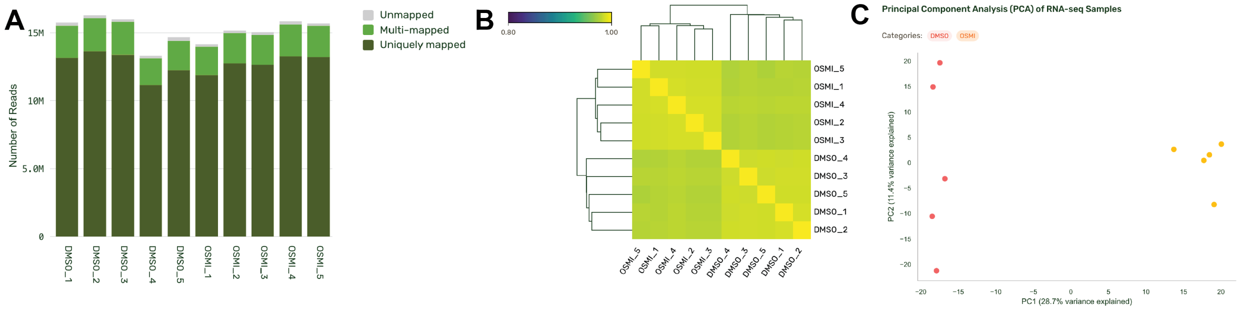

**Figure S1. RNA-Seq Metrics Processed by Plasmidsaurus. A)** Read counts per sample filtered using FastP v0.24.0: poly-X tail trimming, 3' quality-based trimming, minimum quality score of 15 and minimum length requirement of 50 bp. Alignment to human reference genome using STAR aligner v2.8 with non-canonical splice junctions removed. **B)** Sample-sample correlations calculated on normalized counts (TMM, trimmed mean of M-values) using Pearson correlation shown as a heatmap. **C)** Principal component analysis calculated on normalized counts showing clustering within treatment groups.

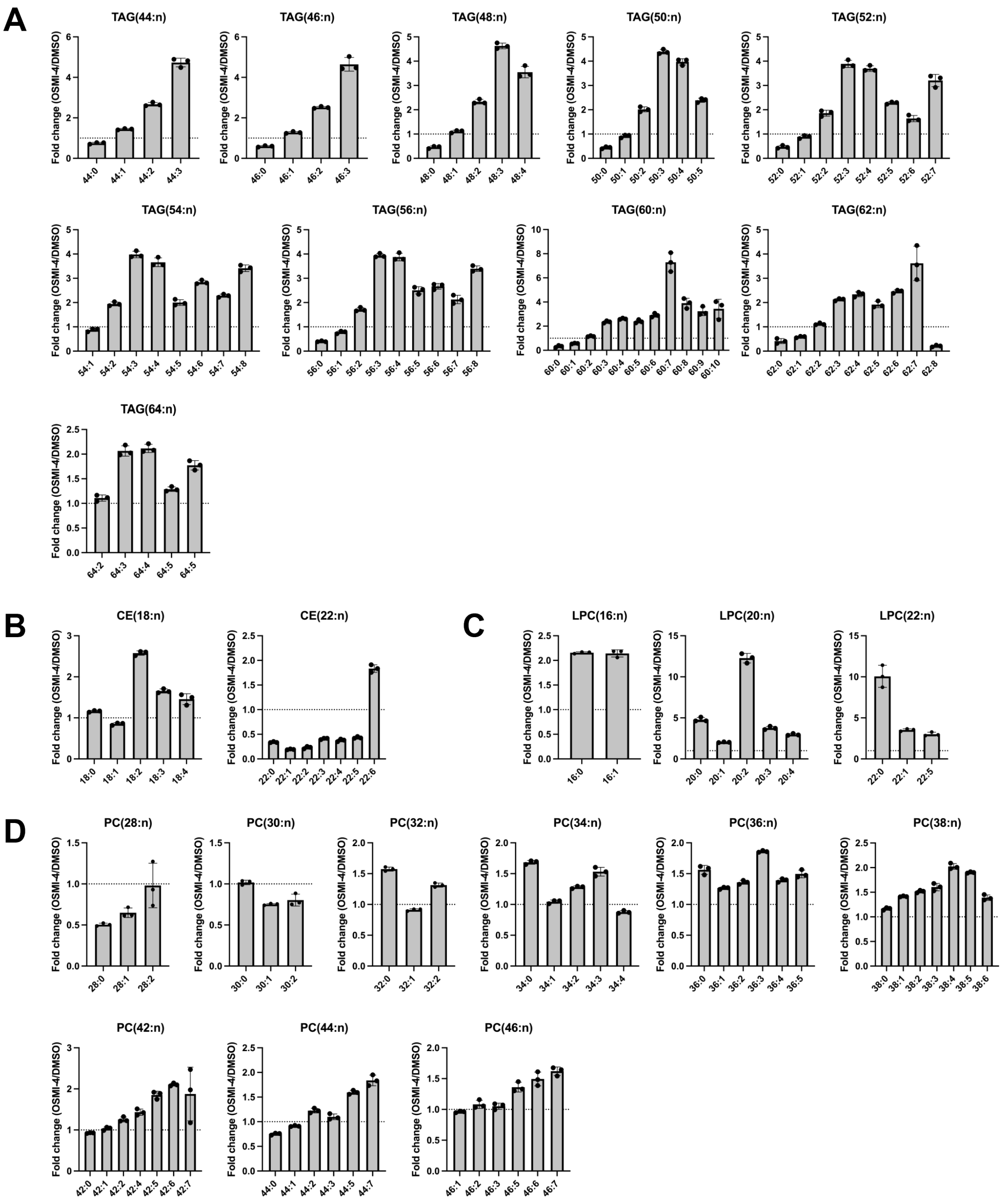

**Figure S2. Quantification of lipidomic profiling data for the remaining specific by lipid class.** Lipidomic profiling was performed with *n*=3 biological replicates. **A)** TAGs **B)** CEs **C)** LPCs **D)** PCs.

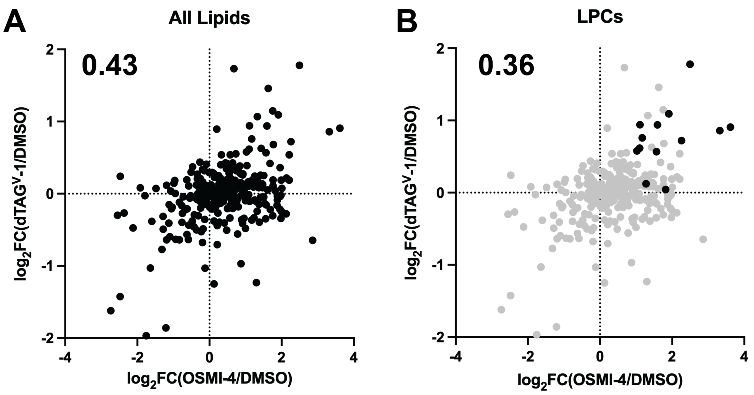

**Figure S3. Plots correlating fold-change in lipid specific abundance between OSMI-4 treatment and dTAG-v1 treatment in OGT-dTAG HCT 116 cells. A)** All lipid species. **B)** Black dots indicate the LPCs, and the grey dots represent all other lipid species. Number displayed on the plots are the Pearson correlation coefficients calculated using the fold changes.

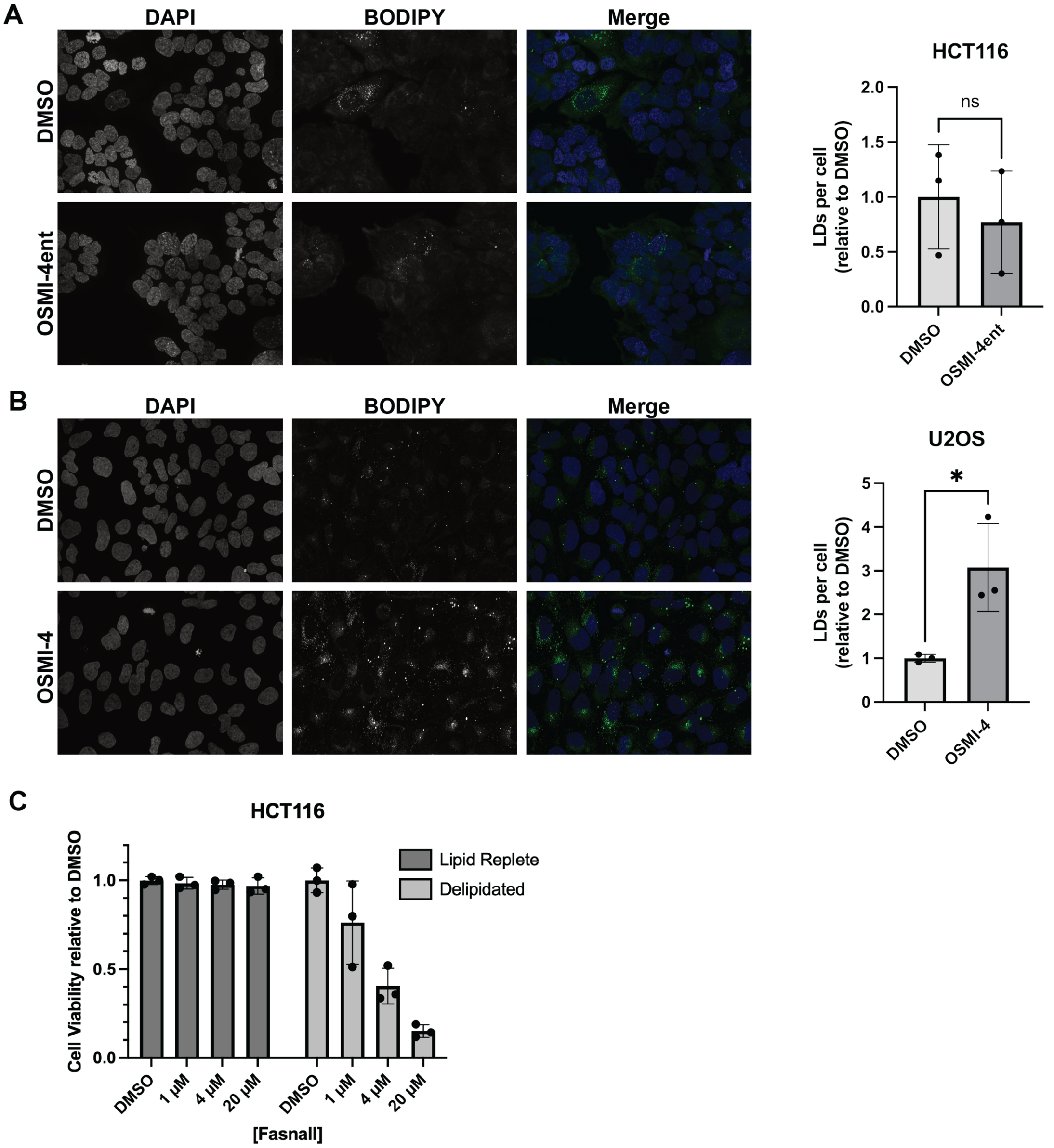

**Figure S4. A)** Representative images of lipid droplets (LDs) and quantification LDs per cell from confocal images of HCT 116 cells treated with either DMSO or 20 µM OSMI-4ent for 24 h. **B)** Representative images and quantification of LDs per cell of confocal images from U2OS cells treated with either DMSO or 20 µM OSMI-4 for 24 h. Analysis was performed with *n*=3 biological replicates. **C)** Dose response to fansall in HCT 116 cells grown in delipidated media and lipid replete media control after 24 hr of treatment. Dose response was performed with n=3 biological replicates. Error bars represent standard deviation.

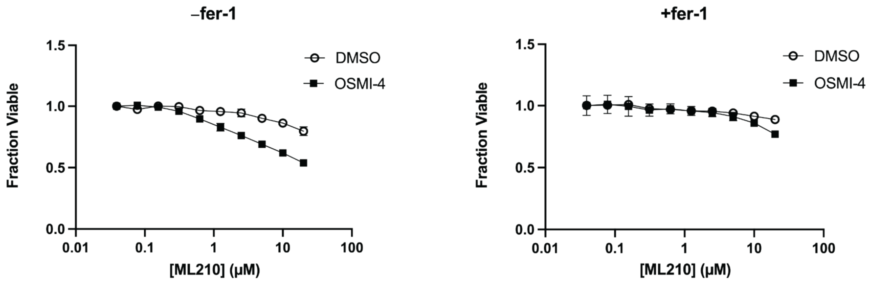

**Figure S5.** Dose response of HCT 116 cells to ML210 with or without 20 μM OSMI-4 and with or without 15 μM ferrostatin-1 after 3 days of co-treatment. Dose response was performed with *n*=3 biological replicates.

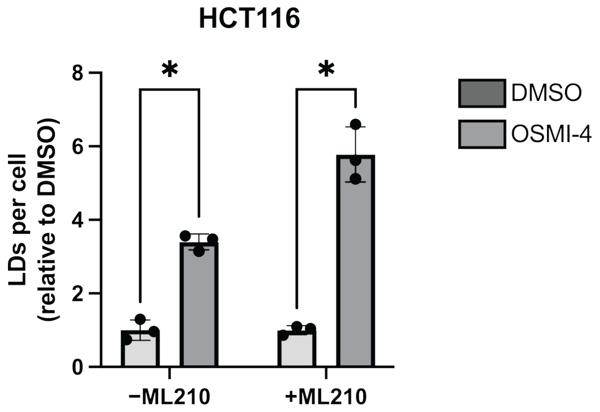

**Figure S6.** Quantification of LDs per cell from confocal images of HCT 116 cells treated with DMSO or 20 μM OSMI-4 for 24 h treatment and co-treatment with 5 μM ML210. Experiments were performed with *n*=3 biological replicates.

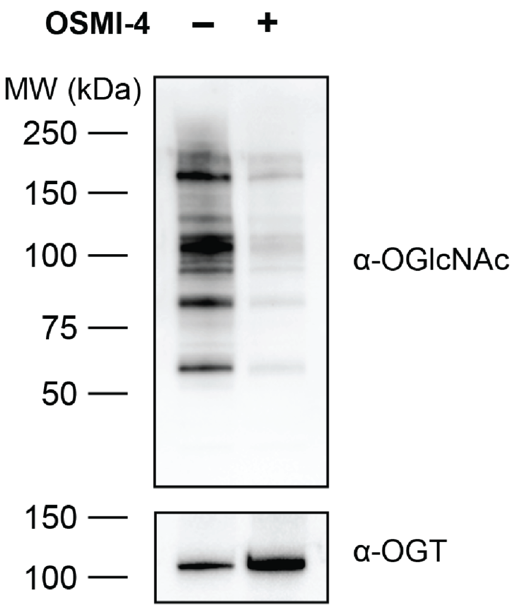

**Figure S7. Western blot of cell lysates from HT-1080 cells.** Cells were treated with DMSO or 20 µM OSMI-4 for 24 h. Blotting against OGT and O-GlcNAc (using RL2 antibody).

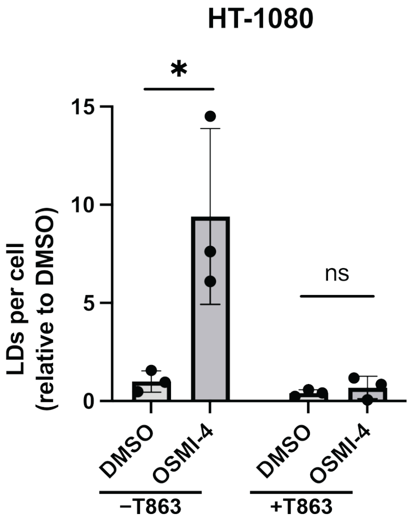

**Figure S8.** Quantification of LDs per cell from confocal images of HT-1080 cells co-treated with 20 µM OSMI-4 and 2 µM T863 for 24 h.

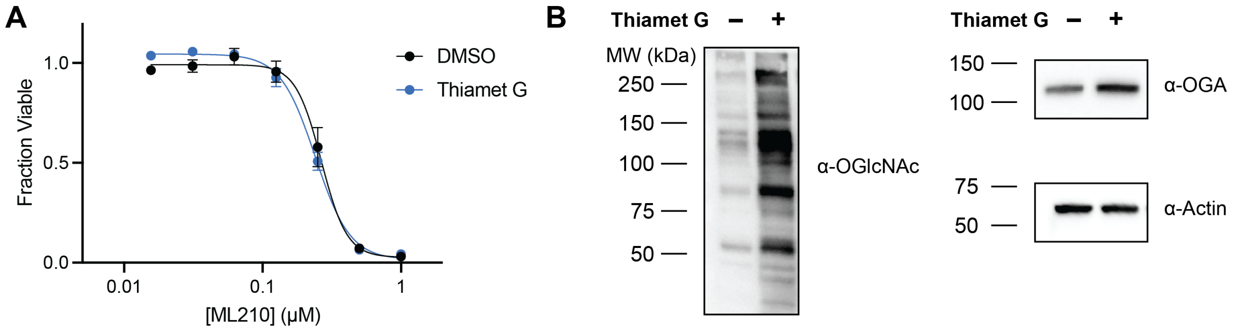

**Figure S9. A)** Dose response of HT 1080 cells to ML210 with or without 10 µM Thiamet G for 48 h. Experiments were performed with *n*=3 biological replicates. **B)** Western blot of cell lysates from HT-1080 cells to ensure sufficient OGA inhibition. Cells were treated with DMSO or 10 µM TMG for 48 h. Blotting against OGA and O-GlcNAc (using RL2 antibody).

### Supplemental Tables

**Table S1.** Upregulated genes from RNA-Seq of HCT 116 cell treated with OSMI vs DMSO control. Hits defined as $log_2FC > 1$, p value < 0.05 and logCPM > 1.

| EnsemblID | $Log_2FC$ (OSMI/DMSO) | logCPM | p Value | gene_name |
|---|---:|---:|---:|---|
| ENSG00000079459 | 1.81914 | 7.1847 | 1.17E-23 | FDFT1 |
| ENSG00000186480 | 2.49584 | 5.96044 | 3.63E-23 | INSIG1 |
| ENSG00000067064 | 1.65918 | 6.08189 | 3.33E-21 | IDI1 |
| ENSG00000099194 | 1.73593 | 7.84647 | 4.30E-21 | SCD |
| ENSG00000167508 | 1.50442 | 6.09358 | 9.92E-21 | MVD |
| ENSG00000169174 | 2.15867 | 4.54705 | 2.13E-20 | PCSK9 |
| ENSG00000149485 | 1.63413 | 5.9359 | 6.38E-20 | FADS1 |
| ENSG00000172893 | 1.22453 | 5.82198 | 1.54E-19 | DHCR7 |
| ENSG00000052802 | 1.36636 | 6.0021 | 1.98E-19 | MSMO1 |
| ENSG00000120437 | 1.08675 | 5.81906 | 1.09E-18 | ACAT2 |
| ENSG00000160285 | 1.44348 | 4.81305 | 1.07E-17 | LSS |
| ENSG00000164211 | 1.64091 | 3.89404 | 2.88E-17 | STARD4 |
| ENSG00000113161 | 1.02681 | 6.44617 | 4.94E-17 | HMGCR |
| ENSG00000134324 | 1.77716 | 3.26805 | 5.95E-17 | LPIN1 |
| ENSG00000130164 | 1.18916 | 5.56383 | 1.05E-16 | LDLR |
| ENSG00000170522 | 1.1813 | 5.11099 | 1.26E-16 | ELOVL6 |
| ENSG00000137440 | 2.44217 | 2.84585 | 3.34E-16 | FGFBP1 |
| ENSG00000122861 | 1.00517 | 5.5487 | 3.87E-16 | PLAU |
| ENSG00000149809 | 1.37144 | 3.9729 | 9.66E-16 | TM7SF2 |
| ENSG00000217555 | 1.01677 | 5.08693 | 1.24E-15 | CKLF |
| ENSG00000110921 | 1.23093 | 4.33296 | 2.54E-15 | MVK |
| ENSG00000196408 | 1.51242 | 3.298 | 4.50E-15 | NOXO1 |
| ENSG00000203778 | 1.20779 | 3.91044 | 5.25E-15 | FAM229B |
| ENSG00000133216 | 1.25694 | 4.02028 | 8.35E-15 | EPHB2 |
| ENSG00000023839 | 1.03566 | 4.32321 | 1.43E-14 | ABCC2 |
| ENSG00000181577 | 2.81919 | 1.61213 | 2.40E-14 | LINC03040 |
| ENSG00000141574 | 1.36361 | 3.42578 | 3.83E-14 | SECTM1 |
| ENSG00000169583 | 1.73518 | 2.57115 | 7.19E-14 | CLIC3 |
| ENSG00000170915 | 2.40715 | 1.90352 | 1.32E-13 | PAQR8 |
| ENSG00000163683 | 1.73209 | 3.53249 | 1.56E-13 | SMIM14 |
| ENSG00000109654 | 1.38889 | 3.76199 | 2.54E-13 | TRIM2 |
| ENSG00000181392 | 1.09284 | 4.05925 | 3.24E-13 | SYNE4 |
| ENSG00000003147 | 1.32829 | 3.14526 | 3.40E-13 | ICA1 |

| | | | | |
|---|---|---|---|---|
| ENSG00000172086 | 1.44737 | 2.78424 | 7.88E-13 | KRCC1 |
| ENSG00000197191 | 1.78895 | 2.21643 | 8.64E-13 | CYSRT1 |
| ENSG00000111671 | 1.11302 | 3.82035 | 1.42E-12 | SPSB2 |
| ENSG00000143643 | 1.33544 | 2.63307 | 1.44E-12 | TTC13 |
| ENSG00000179178 | 1.09434 | 3.19163 | 1.63E-12 | TMEM125 |
| ENSG00000138764 | 1.46616 | 2.56045 | 1.89E-12 | CCNG2 |
| ENSG00000186104 | 1.1764 | 2.85894 | 4.12E-12 | CYP2R1 |
| ENSG00000078114 | 1.17223 | 2.66841 | 5.02E-12 | NEBL |
| ENSG00000074181 | 1.1661 | 2.71081 | 8.99E-12 | NOTCH3 |
| ENSG00000060642 | 1.05765 | 3.51135 | 1.15E-11 | PIGV |
| ENSG00000119242 | 1.29657 | 3.15429 | 1.26E-11 | CCDC92 |
| ENSG00000005001 | 1.2543 | 2.58891 | 1.50E-11 | PRSS22 |
| ENSG00000167861 | 1.19699 | 2.83141 | 1.76E-11 | HID1 |
| ENSG00000049283 | 1.22075 | 2.41882 | 2.41E-11 | EPN3 |
| ENSG00000109107 | 1.12625 | 2.81439 | 2.90E-11 | ALDOC |
| ENSG00000105289 | 1.65177 | 2.40415 | 5.02E-11 | TJP3 |
| ENSG00000140465 | 1.82182 | 1.84703 | 6.05E-11 | CYP1A1 |
| ENSG00000133466 | 1.22068 | 2.25117 | 6.62E-11 | C1QTNF6 |
| ENSG00000108846 | 1.15244 | 3.24343 | 9.54E-11 | ABCC3 |
| ENSG00000166750 | 1.19434 | 2.61941 | 1.03E-10 | SLFN5 |
| ENSG00000118898 | 1.05881 | 3.41109 | 2.53E-10 | PPL |
| ENSG00000214274 | 1.04222 | 3.09063 | 2.70E-10 | ANG |
| ENSG00000147883 | 1.66339 | 1.66209 | 2.95E-10 | CDKN2B |
| ENSG00000068024 | 1.12201 | 2.3116 | 3.12E-10 | HDAC4 |
| ENSG00000168517 | 1.0329 | 2.70967 | 3.27E-10 | HEXIM2 |
| ENSG00000198121 | 1.05815 | 2.86902 | 4.83E-10 | LPAR1 |
| ENSG00000165102 | 1.27129 | 2.97768 | 6.22E-10 | HGSNAT |
| ENSG00000107821 | 1.18441 | 1.94647 | 6.25E-10 | KAZALD1 |
| ENSG00000184574 | 1.12468 | 2.34151 | 7.46E-10 | LPAR5 |
| ENSG00000013563 | 1.15324 | 2.35555 | 8.80E-10 | DNASE1L1 |
| ENSG00000025423 | 1.39149 | 1.3318 | 1.89E-09 | HSD17B6 |
| ENSG00000075240 | 1.14234 | 2.58081 | 2.00E-09 | GRAMD4 |
| ENSG00000163467 | 1.30486 | 2.15168 | 2.08E-09 | TSACC |
| ENSG00000078018 | 1.10344 | 1.99618 | 4.62E-09 | MAP2 |
| ENSG00000187720 | 1.20863 | 2.55469 | 5.50E-09 | THSD4 |
| ENSG00000188177 | 1.19975 | 1.88726 | 5.86E-09 | ZC3H6 |
| ENSG00000153982 | 1.10669 | 2.66486 | 7.62E-09 | GDPD1 |
| ENSG00000258818 | 1.16093 | 1.61215 | 7.69E-09 | RNASE4 |
| ENSG00000165730 | 1.01604 | 2.26174 | 8.68E-09 | STOX1 |

| | | | | |
|---|---|---|---|---|
| ENSG00000259171 | 1.15658 | 1.59287 | 8.96E-09 | |
| ENSG00000205795 | 1.51096 | 1.49651 | 1.36E-08 | CYS1 |
| ENSG00000130517 | 1.01371 | 3.18896 | 1.70E-08 | PGPEP1 |
| ENSG00000102934 | 1.01701 | 1.73358 | 2.10E-08 | PLLP |
| ENSG00000232859 | 1.35761 | 1.44321 | 3.11E-08 | LYRM9 |
| ENSG00000135709 | 1.00484 | 1.9483 | 4.44E-08 | KIAA0513 |
| ENSG00000182612 | 1.26484 | 1.39484 | 6.38E-08 | TSPAN10 |
| ENSG00000118785 | 1.29157 | 1.24643 | 6.95E-08 | SPP1 |
| ENSG00000105982 | 1.09967 | 1.82497 | 7.17E-08 | RNF32 |
| ENSG00000125864 | 1.05761 | 2.21529 | 1.05E-07 | BFSP1 |
| ENSG00000197889 | 1.0828 | 1.39478 | 1.96E-07 | MEIG1 |
| ENSG00000203780 | 1.09127 | 2.17368 | 1.96E-07 | FANK1 |
| ENSG00000071246 | 1.16652 | 1.49654 | 2.19E-07 | VASH1 |
| ENSG00000153132 | 1.19575 | 1.34076 | 2.30E-07 | CLGN |
| ENSG00000171914 | 1.10596 | 1.35356 | 2.50E-07 | TLN2 |
| ENSG00000162620 | 1.33257 | 1.03717 | 3.24E-07 | LRRIQ3 |
| ENSG00000153558 | 1.00998 | 1.57767 | 7.31E-07 | FBXL2 |
| ENSG00000164307 | 1.05616 | 1.52042 | 8.23E-07 | ERAP1 |
| ENSG00000175938 | 1.05222 | 1.18487 | 1.97E-06 | ORAI3 |
| ENSG00000120278 | 1.05273 | 1.03741 | 2.67E-06 | PLEKHG1 |
| ENSG00000188747 | 1.18328 | 1.3622 | 2.93E-06 | NOXA1 |
| ENSG00000156500 | 1.17268 | 1.51984 | 1.09E-05 | PABIR3 |
| ENSG00000100271 | 1.06163 | 1.38049 | 1.29E-05 | TTLL1 |
| ENSG00000117472 | 1.05937 | 1.05833 | 2.24E-05 | TSPAN1 |

**Table S2.** Downregulated genes from RNA-Seq of HCT 116 cell treated with OSMI vs DMSO control. Hits defined as $log_2FC < -1$, p value < 0.05 and logCPM > 1.

| EnsemblID | $Log_2FC$ (OSMI/DMSO) | logCPM | p Value | gene_name |
|---|---|---|---|---|
| ENSG00000128228 | -1.61043 | 7.09685 | 6.72E-22 | SDF2L1 |
| ENSG00000109084 | -1.37323 | 5.99099 | 5.76E-21 | TMEM97 |
| ENSG00000179958 | -1.38158 | 6.94003 | 3.66E-20 | DCTPP1 |
| ENSG00000120738 | -1.92137 | 5.0428 | 5.20E-20 | EGR1 |
| ENSG00000198408 | -1.54337 | 5.25966 | 1.27E-19 | OGA |
| ENSG00000158716 | -1.81936 | 4.43796 | 2.47E-19 | DUSP23 |
| ENSG00000130204 | -1.0559 | 7.07179 | 6.29E-19 | TOMM40 |
| ENSG00000183010 | -1.02997 | 7.04778 | 9.05E-19 | PYCR1 |
| ENSG00000140332 | -1.49573 | 4.02416 | 1.82E-17 | TLE3 |
| ENSG00000114767 | -1.09685 | 5.51032 | 1.85E-17 | RRP9 |
| ENSG00000212724 | -1.64382 | 3.76041 | 3.59E-17 | KRTAP2-3 |
| ENSG00000130475 | -1.30633 | 4.99291 | 4.80E-17 | FCHO1 |
| ENSG00000241360 | -1.54397 | 4.16111 | 1.77E-16 | PDXP |
| ENSG00000120158 | -1.27282 | 4.2205 | 2.39E-16 | RCL1 |
| ENSG00000285304 | -1.44139 | 4.23031 | 3.87E-16 | |
| ENSG00000108439 | -1.04382 | 5.03885 | 9.65E-16 | PNPO |
| ENSG00000099849 | -1.08718 | 4.2646 | 1.37E-15 | RASSF7 |
| ENSG00000178821 | -1.11668 | 4.46345 | 2.11E-15 | TMEM52 |
| ENSG00000226887 | -1.13182 | 4.40237 | 3.41E-15 | ERVMER34-1 |
| ENSG00000063241 | -1.36031 | 4.17905 | 2.01E-14 | ISOC2 |
| ENSG00000114315 | -1.03381 | 4.42738 | 4.04E-14 | HES1 |
| ENSG00000198585 | -1.11089 | 3.51821 | 9.14E-14 | NUDT16 |
| ENSG00000140854 | -1.02683 | 4.21591 | 1.10E-13 | KATNB1 |
| ENSG00000162062 | -1.06908 | 4.2365 | 1.20E-13 | TEDC2 |
| ENSG00000136856 | -1.11088 | 3.78515 | 4.28E-13 | SLC2A8 |
| ENSG00000158050 | -1.03183 | 4.08492 | 8.75E-13 | DUSP2 |
| ENSG00000198435 | -1.1825 | 3.54522 | 1.41E-12 | NRARP |
| ENSG00000102967 | -1.11155 | 3.97141 | 1.42E-12 | DHODH |
| ENSG00000115271 | -1.295 | 3.05449 | 1.75E-12 | GCA |
| ENSG00000141542 | -1.16789 | 3.02419 | 1.87E-12 | RAB40B |
| ENSG00000148175 | -1.02885 | 3.62651 | 2.05E-12 | STOM |
| ENSG00000163376 | -1.54334 | 1.96818 | 4.74E-12 | KBTBD8 |
| ENSG00000204519 | -1.35935 | 2.83592 | 7.30E-12 | ZNF551 |
| ENSG00000106689 | -1.13335 | 2.82207 | 1.37E-11 | LHX2 |
| ENSG00000118263 | -1.43484 | 2.07822 | 1.96E-11 | KLF7 |
| ENSG00000172059 | -1.70682 | 2.29956 | 4.33E-11 | KLF11 |

| | | | | |
|---|---|---|---|---|
| ENSG00000160161 | -1.08607 | 2.81409 | 7.69E-11 | CILP2 |
| ENSG00000123685 | -1.03505 | 2.70477 | 1.18E-10 | BATF3 |
| ENSG00000101400 | -1.03292 | 3.04447 | 2.12E-10 | SNTA1 |
| ENSG00000131845 | -1.29797 | 1.91905 | 2.19E-10 | ZNF304 |
| ENSG00000105649 | -1.67173 | 1.40641 | 2.50E-10 | RAB3A |
| ENSG00000172197 | -1.20567 | 1.9508 | 3.46E-10 | MBOAT1 |
| ENSG00000105875 | -1.37964 | 1.74554 | 3.60E-10 | WDR91 |
| ENSG00000071282 | -1.8529 | 1.24321 | 8.79E-10 | LMCD1 |
| ENSG00000143333 | -1.07316 | 2.22032 | 1.39E-09 | RGS16 |
| ENSG00000119915 | -1.23025 | 2.07107 | 1.77E-09 | ELOVL3 |
| ENSG00000259330 | -1.06415 | 2.79607 | 1.86E-09 | INAFM2 |
| ENSG00000107485 | -1.61805 | 1.09659 | 4.14E-09 | GATA3 |
| ENSG00000166289 | -1.26668 | 2.00929 | 1.00E-08 | PLEKHF1 |
| ENSG00000114737 | -1.23413 | 1.7877 | 3.25E-08 | CISH |
| ENSG00000189221 | -1.1377 | 1.77486 | 3.35E-08 | MAOA |
| ENSG00000175691 | -1.07174 | 1.86399 | 4.82E-08 | ZNF77 |
| ENSG00000160223 | -1.13253 | 1.48003 | 8.37E-08 | ICOSLG |
| ENSG00000005513 | -1.32137 | 1.01176 | 1.57E-07 | SOX8 |
| ENSG00000163623 | -1.22401 | 1.47525 | 1.70E-07 | NKX6-1 |
| ENSG00000160233 | -1.25019 | 1.21538 | 3.21E-07 | LRRC3 |
| ENSG00000071242 | -1.07919 | 1.69191 | 4.58E-07 | RPS6KA2 |
| ENSG00000186523 | -1.02462 | 1.22665 | 4.79E-07 | FAM86B1 |
| ENSG00000183060 | -1.04873 | 1.81312 | 6.23E-07 | LYSMD4 |
| ENSG00000149781 | -1.03537 | 1.07981 | 7.41E-07 | FERMT3 |
| ENSG00000153933 | -1.03989 | 1.3006 | 1.53E-06 | DGKE |
| ENSG00000188493 | -1.05038 | 1.48488 | 2.16E-06 | ACTMAP |
| ENSG00000121207 | -1.07737 | 1.22947 | 2.71E-06 | LRAT |
| ENSG00000100918 | -1.02487 | 1.01108 | 9.82E-06 | REC8 |
| ENSG00000155367 | -1.10949 | 1.29741 | 1.08E-05 | PPM1J |
| ENSG00000174804 | -1.03681 | 1.225 | 1.68E-05 | FZD4 |

**Table S3.** Results of Gene Set Enrichment Analysis run with "Reactome_Pathways_2024" gene set using GSEApy and Enrichr shown in Figure 1C.

| Term | P-value | Adjusted P-value | $-\log_{10}$ adjusted P-value | Genes |
|---|---|---|---|---|
| Cholesterol Biosynthesis | 6.88E-15 | 2.55E-12 | 11.59 | IDI1; MVK; MSMO1; MVD; HMGCR; DHCR7; LSS; ACAT2; TM7SF2; FDFT1 |
| Metabolism of Steroids | 1.46E-13 | 2.71E-11 | 10.57 | ABCC3; IDI1; STARD4; MVK; INSIG1; MSMO1; ELOVL6; HMGCR; LSS; ACAT2; TM7SF2; SCD; CYP2R1; MVD; DHCR7; FDFT1 |
| Regulation of Cholesterol Biosynthesis by SREBP (SREBF) | 7.60E-13 | 9.40E-11 | 10.03 | IDI1; MVK; SCD; INSIG1; ELOVL6; MVD; HMGCR; DHCR7; LSS; TM7SF2; FDFT1 |
| Activation of Gene Expression by SREBF (SREBP) | 1.40E-12 | 1.30E-10 | 9.89 | IDI1; MVK; SCD; ELOVL6; MVD; HMGCR; DHCR7; LSS; TM7SF2; FDFT1 |
| Metabolism of Lipids | 5.00E-08 | 3.71E-06 | 5.43 | ABCC3; IDI1; STARD4; MVK; INSIG1; ELOVL3; GDPD1; MBOAT1; MSMO1; ELOVL6; HMGCR; LSS; ACAT2; TM7SF2; DUSP23; SCD; CYP2R1; CYP1A1; MVD; DHCR7; LPIN1; FADS1; FDFT1 |
| Metabolism | 5.61E-05 | 3.47E-03 | 2.46 | IDI1; MVK; MAOA; INSIG1; GDPD1; MSMO1; HMGCR; ACAT2; TM7SF2; HYAL1; CYP2R1; PNPO; LRAT; NUDT16; LDLR; FADS1; FDFT1; ABCC3; HGSNAT; STARD4; ABCC2; ELOVL3; MBOAT1; PYCR1; ELOVL6; LSS; DHODH; DUSP23; SCD; CYP1A1; ALDOC; MVD; DHCR7; LPIN1; DCTPP1 |
| Linoleic Acid (LA) Metabolism | 1.77E-03 | 9.37E-02 | 1.03 | ELOVL3; FADS1 |

**Table S4.** Fold change scores for lipid classes detected in lipidomics. Fold change scores are calculated as a geometric mean of the fold change values (OSMI-4 treatment compared to DMSO) for all species within that class.

| **Lipid Class** | **FC score** |
|---|---:|
| Cer | 0.58 |
| CE | 0.56 |
| DAG | 0.96 |
| LPC | 3.47 |
| LPE | 1.59 |
| LPI | 1.4 |
| LPS | 1.64 |
| PC | 1.29 |
| PE | 1.21 |
| PG | 1.37 |
| PI | 1.02 |
| PS | 1.03 |
| SM | 1.87 |
| TAG | 1.78 |

**Table S5.** Saturation scores for lipid classes detected in lipidomics. Scores were calculated based on a formula that takes into account the fold-change (OSMI-4 treatment compared to DMSO) for each lipid species in that class and the number of acyl chains for lipids in that class (see Methods).

| **Lipid Class** | **Sat. Score** |
|---|---:|
| Cer | -0.10 |
| CE | 0.51 |
| DAG | 0.14 |
| LPC | -0.20 |
| LPE | 0.10 |
| LPI | -0.10 |
| LPS | 0.07 |
| PC | 0.13 |
| PE | -0.04 |
| PG | 0.04 |
| PI | 0.03 |
| PS | -0.02 |
| SM | -0.05 |
| TAG | 0.17 |

**Table S6.** Fold change scores for lipid classes detected in lipidomics. Fold change scores are calculated as a geometric mean of the fold change values (dTAG-v1 treatment compared to DMSO) for all species within that class.

| **Lipid Class** | **FC score** |
|---|---:|
| Cer | 1 |
| CE | 0.67 |
| DAG | 1 |
| LPC | 1.71 |
| LPE | 1.04 |
| LPI | 0.63 |
| LPS | 1.15 |
| PC | 1.12 |
| PE | 1.08 |
| PI | 1.05 |
| PS | 1.03 |
| SM | 0.95 |
| TAG | 0.91 |

**Table S7.** Detection parameters and MS intensity values for all detectable lipid species in HCT 116 cells treated with DMSO vs. OSMI-4 and DMSO vs. dTAG-v1.

See excel file.
